# Intravenous administration of BCG protects mice against lethal SARS-CoV-2 challenge

**DOI:** 10.1101/2021.08.30.458273

**Authors:** Kerry L. Hilligan, Sivaranjani Namasivayam, Chad S. Clancy, Danielle O’Mard, Sandra D. Oland, Shelly J. Robertson, Paul J. Baker, Ehydel Castro, Nicole L. Garza, Bernard A. P. Lafont, Reed Johnson, Franca Ronchese, Katrin D. Mayer-Barber, Sonja M. Best, Alan Sher

## Abstract

Early events in the host response to SARS-CoV-2 are thought to play a major role in determining disease severity. During pulmonary infection, the virus encounters both myeloid and epithelioid lineage cells that can either support or restrict pathogen replication as well as respond with host protective versus detrimental mediators. In addition to providing partial protection against pediatric tuberculosis, vaccination with bacille Calmette-Guérin (BCG) has been reported to confer non-specific resistance to unrelated pulmonary pathogens, a phenomenon attributed to the induction of long-lasting alterations within the myeloid cell compartment. Here we demonstrate that prior intravenous, but not subcutaneous, administration of BCG protects human-ACE2 transgenic mice against lethal challenge with SARS-CoV-2 and results in reduced viral loads in non-transgenic animals infected with an alpha variant. The observed increase in host resistance was associated with reductions in SARS-CoV-2-induced tissue pathology, inflammatory cell recruitment and cytokine production that multivariate analysis revealed to be only partially related to diminished viral load. We propose that this protection stems from BCG-induced alterations in the composition and function of the pulmonary cellular compartment that impact the innate response to the virus and the ensuing immunopathology.

## Introduction

COVID-19 is a pneumonic disease caused by the newly emerged coronavirus, severe acute respiratory syndrome coronavirus-2 (SARS-CoV-2). A wide spectrum of disease severity is observed in humans, with some patients remaining entirely asymptomatic and others progressing to severe respiratory disease, multiple organ failure and death (Hu et al., 2021). The symptoms and pathology associated with severe forms of COVID-19 are thought to be driven by an overzealous and sustained innate immune response, with disease severity positively correlated with high levels of pro-inflammatory cytokines, NLRP3 inflammasome assembly and myeloid cell activation (Del Valle et al., 2020; Delorey et al., 2021; Melms et al., 2021; Thwaites et al., 2021; Vora et al., 2021).

Type I interferons (IFN-I) are important mediators of anti-viral responses and sub-optimal or delayed IFN-I responses are associated with more severe forms of COVID-19 (Hadjadj et al., 2020; O’Brien et al., 2020; Zhang et al., 2020). However, prolonged production of IFN-I, IFNλ and other innate cytokines, such as IL-6 and TNFα, can contribute to pulmonary pathology through induction of cell death pathways, recruitment of pro-inflammatory cells and disruption of epithelial repair, as well as to vascular damage through pleiotropic effects on endothelial, vascular smooth muscle cells and infiltrating leukocytes (Channappanavar et al., 2016; Israelow et al., 2020; Karki et al., 2021; Major et al., 2020; Ragab et al., 2020; Thacker et al., 2012). Induction of early innate cytokines followed by a tempering of this response may therefore be important determinants of disease outcome. Indeed, prior pro-inflammatory events can be protective against subsequent unrelated infections through pre-existing IFN priming or the recruitment of anti-microbial monocyte-derived cells into the airways (Aegerter et al., 2020; Cheemarla et al., 2021). CCR2-dependent cells, such as monocytes and type-2 dendritic cells, support early viral control in SARS-CoV-2 infection models, with CCR2-KO mice displaying higher viral titers in the lungs 4 days-post infection (Vanderheiden et al., 2021).

Bacille Calmette-Guérin (BCG) is a live attenuated vaccine that is widely used to prevent disseminated tuberculosis in infants and children (Zwerling et al., 2011). In addition to partially protecting against pediatric tuberculosis, BCG vaccination is associated with beneficial, non-specific effects including lower “all-cause” mortality in infants (Biering-Sorensen et al., 2017), reduced viremia following a yellow fever vaccine challenge in adults (Arts et al., 2018), and reduced risk of respiratory infections in the elderly (Giamarellos-Bourboulis et al., 2020). BCG has also proved successful in stimulating anti-tumor immune responses and is an effective treatment for some forms of bladder cancer (Pettenati and Ingersoll, 2018). These non-specific effects of BCG have been attributed to epigenetic and metabolic re-programming of the innate immune system (Arts et al., 2016; Kleinnijenhuis et al., 2012), as well as the re-direction of haematopoiesis towards the rapid generation of protective myeloid subsets (Brook et al., 2020; Cirovic et al., 2020; Kaufmann et al., 2018). In experimental models this is particularly evident when BCG is administered by the intravenous (iv) as opposed to more conventional subcutaneous (sc) route (Kaufmann et al., 2018). In this regard, the iv route of BCG inoculation has also been shown to provide superior protection against *Mycobacterium tuberculosis* infection in non-human primates (Darrah et al., 2020; Sharpe et al., 2016).

Because of its well-described association with non-specific innate protection, BCG administration has been suggested as a possible prophylactic measure for the prevention of SARS-CoV-2 infection and disease (O’Neill and Netea, 2020). Indeed, a number of ecological studies have suggested an association of prior BCG vaccination with lower incidence of COVID-19 disease (Escobar et al., 2020; Rivas et al., 2021). This concept, which remains controversial, is currently being formally tested in a number of clinical trials (Junqueira-Kipnis et al., 2020; Tsilika et al., 2021). In addition, recent pre-clinical work has demonstrated the use of BCG as an adjuvant to boost SARS-CoV-2 specific vaccine-induced protection (Counoupas et al., 2021).

Here we have systematically evaluated the effects of prior BCG inoculation on SARS-CoV-2 pathogenesis in two murine experimental models. The first model employs K18-hACE2 mice that are highly susceptible to lethal infection due to their expression of a transgene for the human ACE2 receptor under control of the keratin-18 promoter (McCray et al., 2007). The second model involves challenge with an alpha (B.1.1.7) variant that is able to productively infect wildtype, non-transgenic animals. We demonstrate that iv delivery of BCG confers a high level of protection against SARS-CoV-2 in both models. The observed BCG-induced resistance is associated with reduced viral titers and pro-inflammatory cytokine production as well as decreased pulmonary pathology. We propose that iv administration of BCG limits the course of SARS-CoV-2 infection through its targeting of innate immune pathways and thus may be a useful platform for identifying early immunologic events affecting the outcome of this disease.

## Results and Discussion

### Intravenous BCG protects K18-hACE2 animals from lethal SARS-CoV-2 challenge

Non-specific BCG-induced protection is reported to act by promoting the generation of primed innate immune subsets, which is achieved most efficiently if BCG is administered intravenously (iv) as opposed to the more conventional subcutaneous (sc) route (Kaufmann et al., 2018). We therefore hypothesized that if BCG can promote host resistance to SARS-CoV-2, iv administration would be more effective than BCG delivered by alternative routes. To test this, K18-hACE2 transgenic mice were inoculated with BCG either sc or iv and then challenged with a lethal dose of the USA-WA1/2020 (WA) SARS-CoV-2 isolate 42 days later (**Fig1A**). Control animals that received PBS iv and mice given BCG sc underwent profound weight loss following SARS-CoV-2 challenge and exhibited clinical signs of severe disease (ruffled fur, hunched posture, lethargy), progressing to a moribund state. In contrast, animals inoculated with BCG iv 42 days prior to SARS-CoV-2 challenge maintained their body weight and displayed minimal signs of disease as assessed by a blinded observer (**Fig1B-C**). Accordingly, most animals in the PBS iv and BCG sc groups succumbed to infection by 12 days post infection (dpi), whereas BCG iv mice were largely protected, with 85% of animals surviving long-term (**Fig1D**). The few surviving control animals recovered their body weight by 15dpi and, as expected, were fully protected upon SARS-CoV-2 re-challenge (data not shown). These data demonstrate that BCG can trigger potent host resistance against SARS-CoV-2 lethality and that the iv route of BCG delivery is critical for determining the level of protection observed.

**Fig 1:**
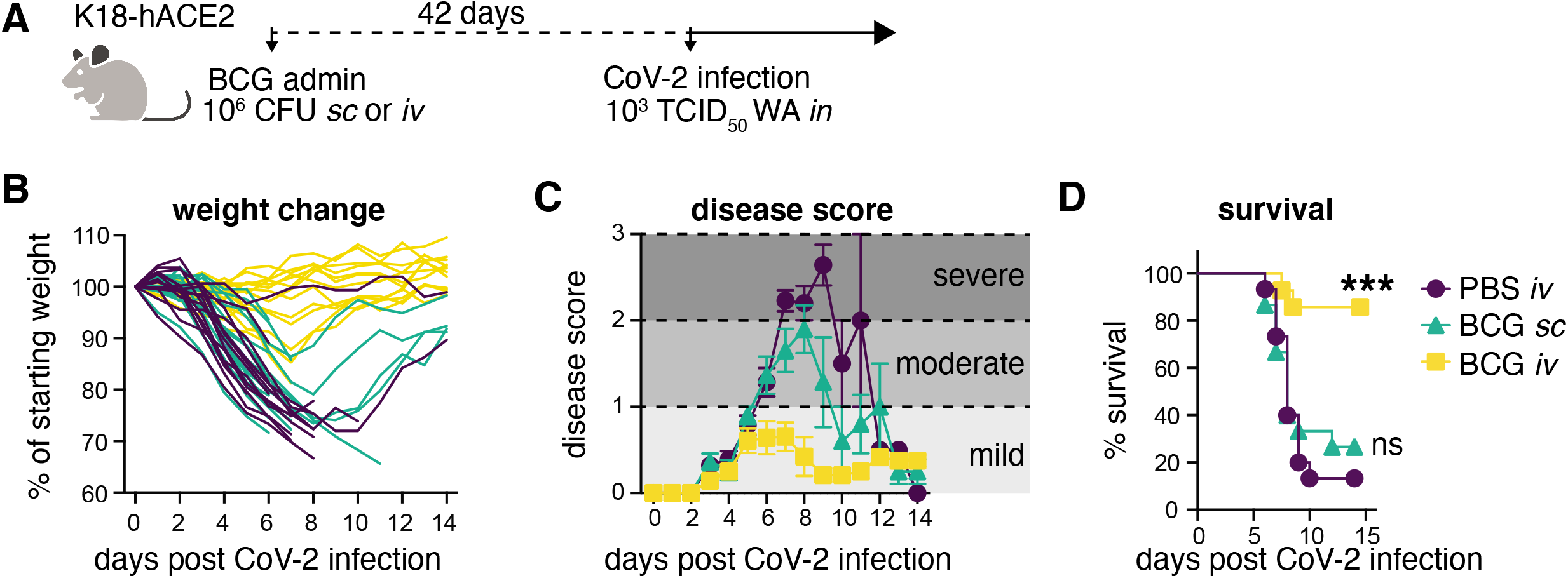
Prior *iv* BCG administration protects K18-hACE2 mice from lethal SARS-CoV-2 challenge. **(A-D)** K18-hACE2 mice were inoculated with 10^6^ CFU BCG Pasteur by subcutaneous (*sc*) or intravenous (*iv*) injection. Control animals received the same volume of PBS *iv*. At 42 days post BCG administration, mice were infected with 10^3^ TCID_50_ SARS-CoV-2 (WA1/2020) by intranasal instillation. **(A)** Schematic of experimental design. **(B)** Weight change following SARS-CoV-2 infection shown as percent of starting weight. **(C)** Animals were scored on a scale of 0-3 by a blinded observer, with 0 referring to no observable signs of disease and 3 referring to a moribund state requiring euthanasia. Scoring criteria are outlined in the Methods. **(D)** Kaplan-Meier curve of animal survival following SARS-CoV-2 challenge. Statistical significance was assessed by Mantel-Cox test. ns *p*>0.05; \*\*\*\**p*<0.0001. Mean±SEM is shown. Data are pooled from 3 independent experiments each with 4-5 mice per group.

### BCG reduces SARS-CoV-2 viral loads in the lungs of K18-hACE2 and wildtype (non-transgenic) mice

To determine whether BCG-induced protection against SARS-CoV-2 correlates with reduced viral loads within tissues, we harvested nasal turbinates, lung and brain from BCG immunized mice 5dpi with SARS-CoV-2 (**Fig2A**). Viral loads were measured by the level of genomic viral RNA specific for the E gene (gRNA) by RT-qPCR and corroborated by tissue culture infectious dose 50 (TCID_50_) assay for lung and brain samples. gRNA was detectable in the nasal turbinates of only half of the animals assessed, independent of treatment group. Viral loads in the turbinates did not differ between BCG-immunized and PBS-treated controls among the animals with detectable viral RNA (**Fig2B**). In the lung, gRNA was detectable in all samples tested and was statistically lower (by 2 logs) in animals that received BCG iv prior to SARS-CoV-2 challenge compared to PBS controls **(Fig2C)**. Assessment of the same lung homogenate samples by TCID_50_ assay revealed that the level of infectious virus was indeed lower in BCG iv mice compared to the PBS iv treatment group, with 50% of samples obtained from BCG iv-treated animals having no measurable infectious virus **(Fig2C**). Expression of the hACE2 transgene was consistent across treatment groups, suggesting that the lower infectivity was not due to BCG induced alterations in transcription of the transgene or selective loss of ACE2 expressing host cells (**FigS1**). These data demonstrate that SARS-CoV-2 productively infects and replicates in K18-hACE2 animals regardless of BCG immunization status, but that viral replication is reduced more effectively in mice previously inoculated with BCG by the iv route.

**Fig 2:**
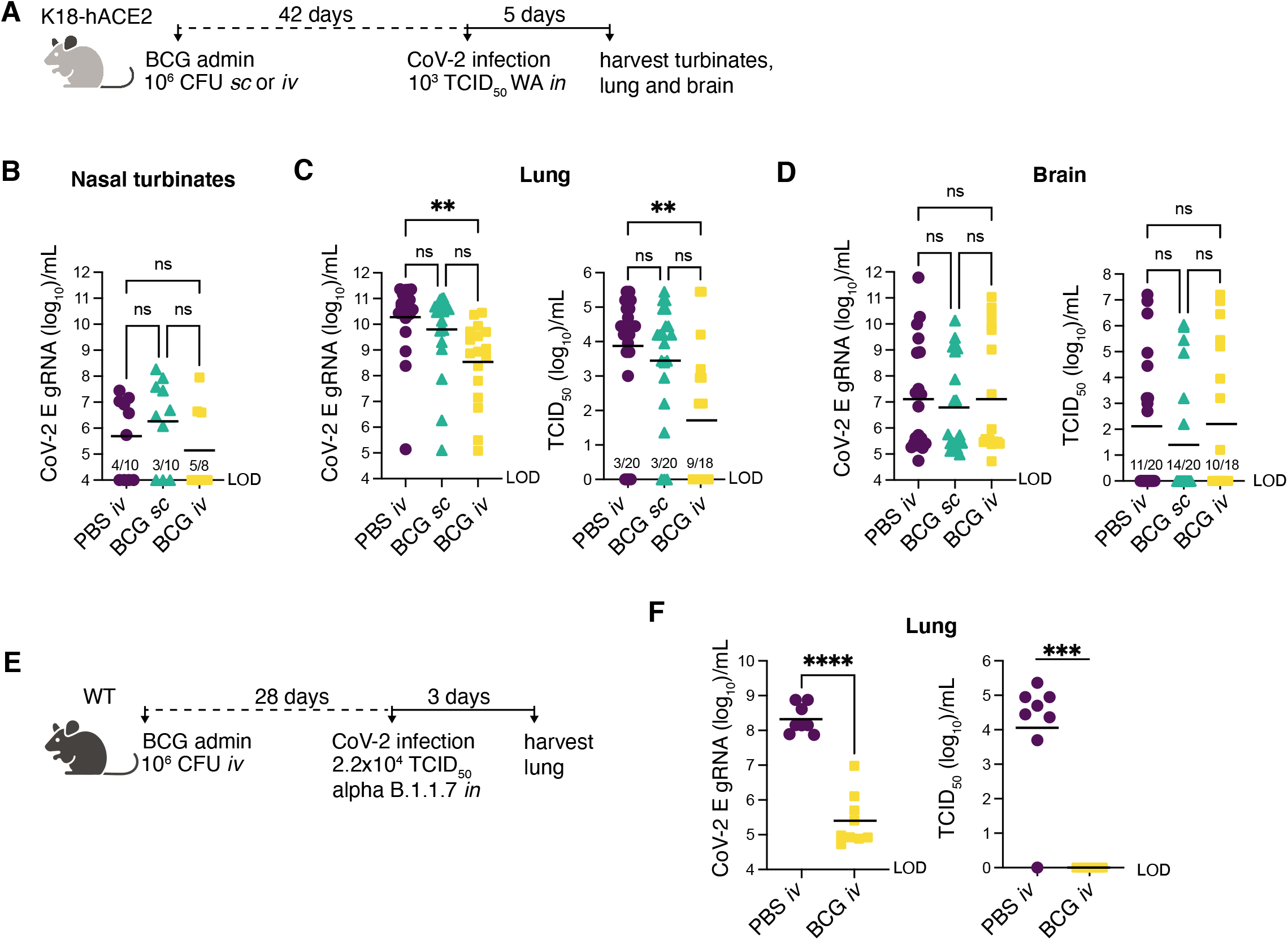
Prior *iv* BCG administration reduces viral loads in the lungs of K18-hACE2 and WT mice. **(A-D)** K18-hACE2 mice were inoculated with 10^6^ CFU BCG Pasteur by subcutaneous (*sc*) or intravenous (*iv*) injection. Control animals received the same volume of PBS *iv*. At 42 days post BCG administration, mice were infected with 10^3^ TCID_50_ SARS-CoV-2 (WA1/2020) by intranasal instillation. Nasal turbinates **(B)**, lungs **(C)** and brain **(D)** were collected 5 days after viral challenge and assessed for viral load by qPCR and TCID_50_ assay. Statistical significance was assessed by Kruskal-Wallis test with Dunn’s posttest. ns *p*>0.05; \*\**p*<0.01. Data are pooled from 3 independent experiments each with 4-5 mice per group. **(E-F)** C57BL/6 (WT) mice were iv injected with 10^6^ CFU BCG Pasteur or PBS. At 28 days post BCG administration, mice were infected with 2.2×10^4^ TCID_50_ SARS-CoV-2 (alpha B.1.1.7) by intranasal instillation. Lungs were collected 3 days after viral challenge and homogenates assessed for viral load by qPCR and TCID_50_ assay **(F)**. Statistical significance was determined by a Mann-Whitney test. \*\*\**p*<0.001; \*\*\*\**p*<0.0001. Data are pooled from 2 independent experiments each with 3-5 mice per group. Graphs show geometric mean and individual values for each animal. LOD (limit of detection).

K18-hACE2 mice are well documented to support neurotropism of SARS-CoV and SARS-CoV-2, which could contribute to mortality in this model (McCray et al., 2007; Yinda et al., 2021). To determine whether the enhanced survival of K18-hACE2 animals conferred by iv BCG was associated with reduced viral loads within the central nervous system (CNS), we assessed gRNA and TCID_50_ titer in brain homogenate 5dpi with SARS-CoV-2. High viral titers (>10^3^ TCID_50_/mL) were observed in a few animals from each treatment group, but there was no difference in viral RNA or infectious virus across the experimental groups (**Fig2D**), arguing against the possibility that iv BCG improves host resistance against SARS-CoV-2 lethality by modulating viral dissemination to the CNS.

Given the artificial expression and tissue distribution of SARS-CoV-2 permissive receptors in K18-hACE2 transgenic mice and the atypical CNS dissemination observed in this model, we tested the efficacy of iv BCG against SARS-CoV-2 in an alternate non-transgenic mouse model. Wildtype mice do not support infection and replication by the originally identified SARS-CoV-2 strains (including the WA isolate) due to incompatibility between the murine ACE2 receptor and the viral spike protein (Zhou et al., 2020). However, recently identified SARS-CoV-2 variants, including the alpha B.1.1.7 variant, have acquired mutations in the spike protein that allows for more efficient engagement of murine ACE2 for viral entry (Montagutelli et al., 2021). We confirmed that wildtype B6 mice could indeed support replication of the B.1.1.7 variant and that this viral expansion was dramatically impaired if the animals had received iv BCG prior to challenge (**Fig2E-F**) thus supporting our findings in the K18-hACE2 mouse model.

Together, these data demonstrated that iv BCG promotes control of SARS-CoV-2 replication in two mouse models. The high level of gRNA copies present in the lungs of iv BCG mice suggested that these animals were productively infected with SARS-CoV-2 but were more efficient at controlling viral replication and infectivity, evidenced by lower infectious viral titers recovered from the lungs. Nevertheless, it was unclear whether the observed level of viral control could account for the striking protection from SARS-CoV-2 driven mortality conferred by iv BCG seen in K18-hACE2 mice. To further examine this issue, we performed a more detailed analysis of the pulmonary immune landscape to identify correlates of protection.

### Prior inoculation with iv BCG reduces SARS-CoV-2 associated pulmonary pathology, immune cell infiltration and chemokine production

Lethal COVID-19 in humans is associated with the accumulation of pro-inflammatory myeloid cells within the lung tissue and airways (Delorey et al., 2021; Melms et al., 2021). Similar associations have been made in murine models that can sustain a productive SARS-CoV-2 infection, including the K18-hACE2 model (Winkler et al., 2020). To identify correlates of iv BCG-induced protection in SARS-CoV-2 infected K18-hACE2 mice, we used histopathology, flow cytometric analysis and cytokine multiplexing to assess the cellular composition and inflammatory environment of lung tissue from BCG iv, BCG sc and PBS iv treated animals 5 days after SARS-CoV-2 challenge (**Fig3A**). We also analyzed the lungs of non-transgenic littermate controls (referred to as wildtype, WT hereafter) inoculated with BCG or PBS and challenged with SARS-CoV-2 WA isolate. As WT animals cannot be productively infected with the WA strain of SARS-CoV-2, these experimental groups allowed us to separate out BCG-induced responses from those driven by SARS-CoV-2 infection.

**Fig 3:**
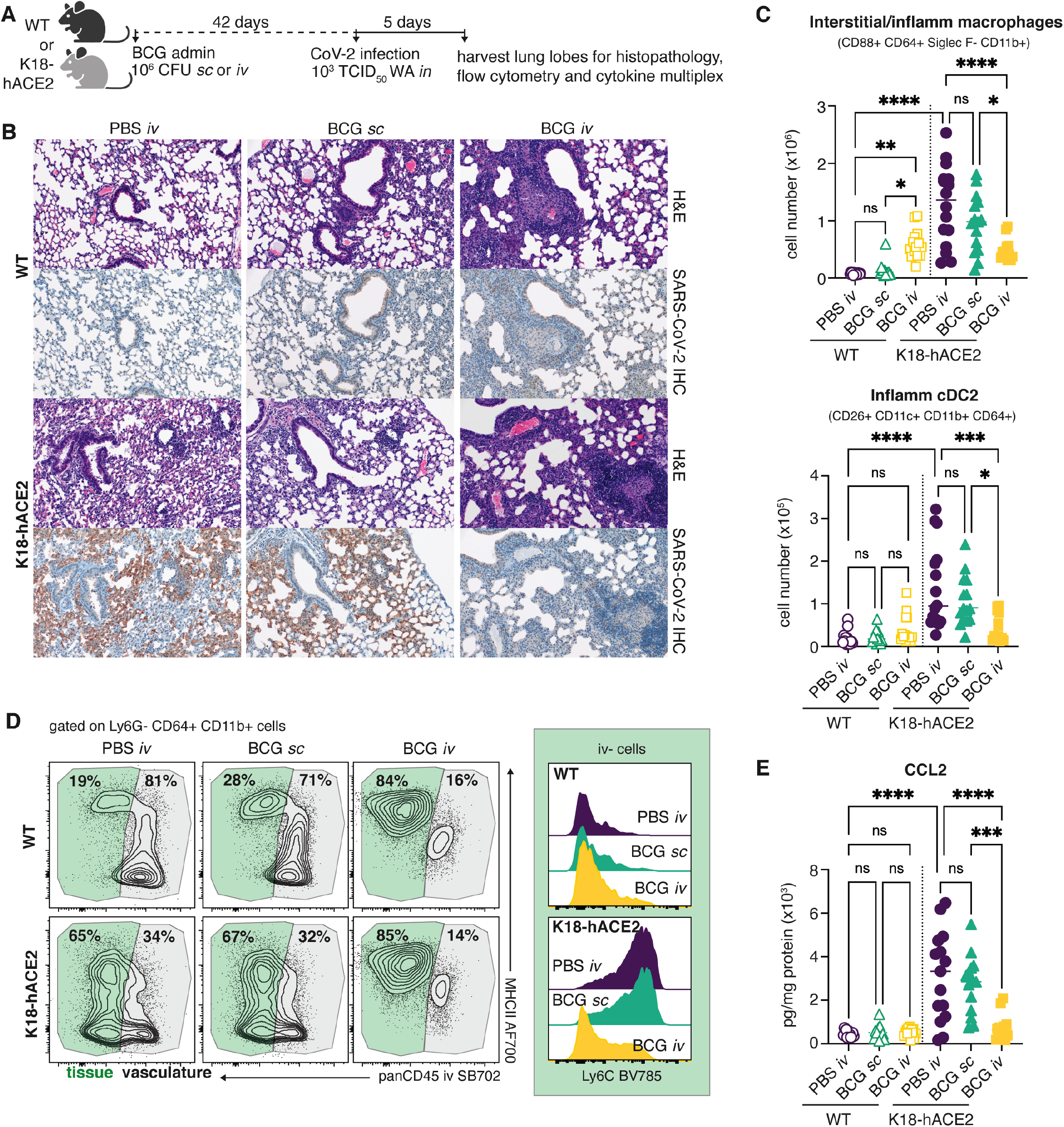
SARS-CoV-2 driven inflammatory cellular responses are dampened by prior *iv* BCG inoculation. **(A-E)** K18-hACE2 mice or non-transgenic littermate controls (WT) were inoculated with 10^6^ CFU BCG Pasteur by subcutaneous (*sc*) or intravenous (*iv*) injection. Control animals received the same volume of PBS *iv*. At 42 days post BCG administration, mice were infected with 10^3^ TCID_50_ SARS-CoV-2 (WA1/2020) by intranasal instillation. Lungs were collected 5 days after viral challenge and assessed by histopathology, flow cytometry and cytokine multiplexing. **(B)** Representative images of hematoxylin and eosin (H&E) stained lung tissue sections (top panels) and sequential sections probed with an anti-CoV-2 nucleoprotein antibody (lower panels). **(C)** Number of interstitial/inflammatory macrophages and inflammatory cDC2. Gating is shown in Supplementary Fig 2. **(D)** Contour plots depict MHCII expression by tissue resident (iv stain negative; green) and vascular (iv stain positive; gray) CD64+ CD11b+ cells in the lung (left). Histograms show Ly6C expression by iv-CD64+ CD11b+ cells (right). Data are concatenated from 4-5 mice per group and are representative of 3 independent experiments. **(E)** CCL2 levels in lung tissue homogenate, standardized to total protein concentration. Statistical significance was assessed by One-Way ANOVA with Tukey post-test. ns *p*>0.05; \**p*<0.05; \*\**p*<0.01; \*\*\**p*<0.001, \*\*\*\**p*<0.0001. Unless otherwise stated, data are pooled from 3 independent experiments each with 4-5 mice per group.

Lungs from WT mice inoculated with BCG iv had moderate pulmonary pathology consisting of discrete, organized granuloma formation within alveolar septa, interstitial pneumonia, influx of foamy macrophages within alveolar spaces and type II pneumocyte hyperplasia in 100% of mice examined. Granulomas were frequently perivascular in distribution, although some appeared to occlude terminal airways. As expected, histopathology did not reveal any lesions consistent with coronaviral respiratory disease in WT animals (**Fig3B**).

In contrast, lesions consistent with respiratory coronavirus infection were observed in 100% of non-BCG and BCG sc inoculated K18-hACE2 mice infected with SARS-CoV-2. Lungs from these animals displayed moderate interstitial pneumonia and leukocyte influx into adjacent alveolar spaces. Bronchiolar disease was absent in all evaluated animals. Consistent with the histopathologic changes observed in the WT groups, interstitial pneumonia with influx of lymphocytes and macrophages was observed in both the BCG iv and sc vaccinated K18-hACE2 mice; however, the lesions associated with SARS-CoV-2 driven pathology were largely absent from mice previously inoculated with BCG iv (**Fig3B**).

Immunohistochemistry for SARS-CoV-2 nucleoprotein was performed to assess the distribution of viral antigen in association with lung pathology. Consistent with our RT-qPCR and TCID_50_ data, SARS-CoV-2 specific immunoreactivity was prominent in sections from virus-challenged K18-hACE2 mice and was localized to numerous, widely dispersed type I and type II pneumocytes as well as macrophages within alveolar spaces. However, antigen distribution was limited in mice inoculated with BCG iv, and was not observed adjacent to pulmonary granulomas (**Fig3B**).

Flow cytometric analysis of digested lung tissue at 5dpi revealed significant upregulation of markers of activation and cell cycle entry (Ki67, CD44, PD1, CD69) by conventional CD8+ and CD4+ T cells, natural killer T (NKT) cells, mucosal associated invariant T (MAIT) cells and gamma-delta T cells in SARS-CoV-2 infected K18-hACE2 mice (**FigS2 and Supplementary Table 1**). Intravenous injection of a fluorescently conjugated panCD45 antibody 3 minutes before euthanasia allowed us to distinguish cells located within the pulmonary vasculature (iv stain+) from cells within the lung parenchyma or airways (iv stain-) (Anderson et al., 2014). SARS-CoV-2 infection promoted recruitment of CD8+ T cells into the lung tissue, evidenced by the striking increase in cells protected from iv staining, as well as a significant increase in Granzyme B+ cells (**FigS3A**). At this timepoint, SARS-CoV-2 (S_539-546_)-specific CD8+ T cells were rare, suggesting that the majority of the observed CD8+ T cell response was non-specific (**FigS3B**). Similarly, SARS-CoV-2 infected mice displayed an increase in parenchymal and Granzyme B+ FoxP3-CD4+ T cells (**FigS3C**). Intravenous BCG administration alone was associated with significant expansion of conventional CD8+, FoxP3-CD4+ and FoxP3+ CD4+ T cells as well as MAIT cells, which did not change following SARS-CoV-2 infection (**FigS3A,C,D,E**). Interestingly, cytotoxic responses were not induced following SARS-CoV-2 infection of mice that had previously received BCG iv, as evidenced by low frequencies of Granzyme B+ T cells and reduced numbers of NK cells compared to infected animals that received PBS iv or BCG sc (**FigS3A,C,E**).

Flow analysis of the myeloid compartment at 5dpi showed significant expansion of CD88+ CD64+ CD11b+ Siglec F-monocyte-derived macrophages in the lungs of SARS-CoV-2 infected animals, consistent with previously published observations (Winkler et al., 2020).We also noted the appearance of a newly described population of inflammatory type-2 dendritic cells (DC2) (CD88-CD26+ CD11c+ MHCII+ XCR1-CD11b+ CD64+) that are regulated by IFN and acquire functional characteristics of macrophages and cross-presenting DC1 (Bosteels et al., 2020) (**Fig3C**). SARS-CoV-2-induced Ly6G-CD64+ CD11b+ cells, containing both DC2 and inflammatory/interstitial subsets, were largely protected from iv CD45 staining, indicating that they were indeed located within the tissue or airways (**Fig3D**). Prior inoculation of WT animals with BCG iv, but not sc, also promoted the accumulation of myeloid cells within the lung, but the phenotype of these cells was that of resident interstitial Ly6C^lo^ MHCII+ macrophages rather than inflammatory Ly6C+ MHCII+/- macrophages. Importantly, productive SARS-CoV-2 infection of animals previously inoculated with BCG iv did not promote further recruitment or expansion of inflammatory macrophages, monocytes or DC2, resulting in an overall lower number of inflammatory cells within the lungs compared to animals infected with SARS-CoV-2 alone (**Fig3C**). In this regard, analysis of lung homogenate revealed SARS-CoV-2 infection induced production of CCL2, a chemokine important for the recruitment of monocytes and DC2 precursors into the lung (Nakano et al., 2017; Serbina and Pamer, 2006). SARS-CoV-2-induced CCL2 production was significantly lower in transgenic mice previously inoculated with BCG iv (**Fig3E**). Together, these observations indicated that SARS-CoV-2 infection results in altered pro-inflammatory and cytotoxic cellular responses in animals previously inoculated iv with BCG compared to control mice given either PBS iv or BCG sc.

### SARS-CoV-2-driven cytokine responses are dampened in iv BCG inoculated mice

Because of the known association of severe COVID-19 with “cytokine storm” (Del Valle et al., 2020; Thwaites et al., 2021), we next assessed the effect of prior iv BCG inoculation on the cytokine milieu of lung tissue from SARS-CoV-2 challenged animals. In keeping with published data sets, SARS-CoV-2 infection drove a marked induction of pro-inflammatory cytokines including IL-6, GM-CSF, IL-12p70 and TNFα in transgenic mice (Winkler et al., 2020), while iv BCG inoculation was associated with elevated IL-12p70 and TNFα production prior to SARS-CoV-2 infection (Wang et al., 1999). Conversely, IL-6 and GM-CSF levels were not induced by iv BCG alone and remained low following SARS-CoV-2 challenge of these animals, resulting in significantly lower levels overall when compared with PBS iv and BCG sc control animals infected with SARS-CoV-2 (**Fig4A**). Consistent with these observations, flow cytometric analysis showed fewer IL-6+ cells in SARS-CoV-2 challenged mice previously inoculated with BCG iv in comparison with PBS iv or BCG sc control animals (**Fig4B**). Of note, we identified CD88+ CD64+ CD11b+ CD11c^mid^ Ly6C^hi^ inflammatory macrophages to be the major source of IL-6 in SARS-CoV-2 challenged mice (**Fig4C**).

**Fig 4:**
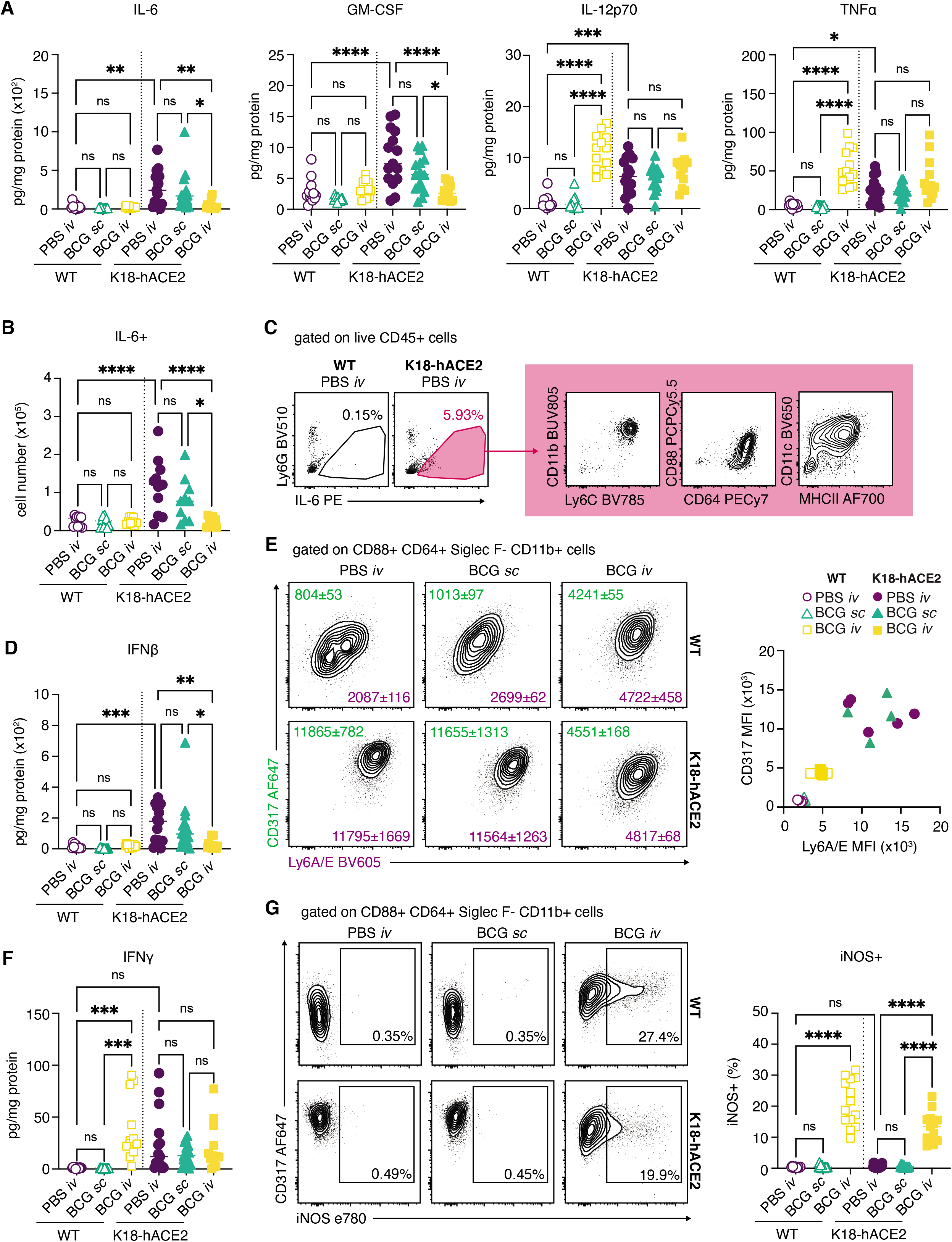
Prior *iv* BCG inoculation alters the inflammatory cytokine milieu of the lung following SARS-CoV-2 challenge. **(A-G)** K18-hACE2 mice or non-transgenic littermate controls (WT) were inoculated with 10^6^ CFU BCG Pasteur by subcutaneous (*sc*) or intravenous (*iv*) routes. Control animals received the same volume of PBS *iv*. At 42 days post BCG administration, mice were infected with 10^3^ TCID_50_ SARS-CoV-2 (WA1/2020) by intranasal instillation. Lungs were collected 5 days after viral challenge and assessed by flow cytometry and cytokine multiplexing. **(A)** IL-6, GM-CSF, IL-12p70 and TNFα levels in lung homogenate, standardized to total protein concentration. **(B)** Number of IL-6+ cells as determined by flow cytometry. **(C)** Gating of IL-6+ cells from total CD45+ cells and surface marker expression of IL-6+ cells from K18-hACE2 PBS iv animals. Plots show data concatenated from 5 animals and are representative of 2 independent experiments. **(D)** IFNβ levels in lung homogenate, standardized to total protein concentration. **(E)** Representative contour plots depict CD317 (green) and Ly6A/E (purple) expression by CD88+ CD64+ Siglec F-CD11b+ macrophages (left). Median fluorescence intensity (MFI) +/- standard error of the mean is indicated on the plots (*n*=4-5). MFI values for individual animals are shown on the right. **(F)** IFNγ levels in lung homogenate, standardized to total protein concentration. **(G)** Representative contour plots depict CD317 and iNOS expression by CD88+ CD64+ Siglec F-CD11b+ macrophages (left). Frequency of iNOS+ macrophages are shown on the right. Statistical significance was assessed by One-Way ANOVA with Tukey post-test. ns *p*>0.05; \**p*<0.05; \*\**p*<0.01; \*\*\**p*<0.001, \*\*\*\**p*<0.0001. Unless otherwise stated, data are pooled from 3 independent experiments each with 4-5 mice per group.

IFN-I are important for control of viral infections but may also contribute to pathology in the context of SARS-CoV-2 infection (Israelow et al., 2020). As expected, SARS-CoV-2 challenge induced IFNβ production in the lungs of PBS iv and BCG sc-treated animals, which was evident by direct measurement of IFNβ in lung homogenate (**Fig4D**) and by the up-regulation of IFN-inducible markers CD317 (BST2, PDCA1) and Ly6A/E (Sca1) on myeloid cells isolated from lung tissue (**Fig4E and Supplementary Table 1**). Importantly, prior iv BCG inoculation suppressed IFNβ induction as well as CD317 and Ly6A/E upregulation following SARS-CoV-2 challenge of transgenic mice. The presence of an ongoing IFN response was suggested by the increased expression of CD317 and Ly6A/E on myeloid cells from WT mice inoculated with BCG iv compared to WT PBS iv and BCG sc-treated animals. While IFNβ levels were low in all WT animals, a robust IFNγ response was observed following iv BCG administration, suggesting that IFNγ production contributes to the induction of IFN-stimulated responses in the lungs of these animals (**Fig4F**). Indeed, a notable population of myeloid cells positive for the IFNγ-inducible enzyme iNOS was detected in the lungs of all iv BCG animals but was absent from PBS iv and BCG sc groups (**Fig4G**).

Together, these data demonstrate that prior iv BCG administration suppresses the SARS-CoV-2-induced inflammatory cytokine response that is known to contribute to tissue damage in both humans and experimental animals infected with SARS-CoV-2 (Del Valle et al., 2020; Israelow et al., 2020; Thwaites et al., 2021). This modulation of pro-inflammatory cytokines may be one of the mechanisms by which iv BCG protects transgenic animals from lethal challenge. Further, the presence of an ongoing BCG-induced IFNγ response in the lungs of iv inoculated animals may support early control of SARS-CoV-2 replication by innate cells, similar to the protective effects observed with IFN-I priming of airway epithelial organoids *in vitro* (Cheemarla et al., 2021).

### The suppression of SARS-CoV-2-induced inflammatory response modules in iv BCG inoculated mice is unrelated to diminished viral loads

To more directly identify correlates of iv BCG-induced protection from SARS-CoV-2 associated disease, we performed a multivariate analysis incorporating weight change, viral loads, flow cytometric parameters and cytokine levels in all of the individual animals studied. A principal component analysis revealed overlapping signatures between PBS iv and BCG sc mice, whereas BCG iv inoculated animals clustered separately (**Fig5A**). SARS-CoV-2 induced changes were evident for PBS iv and BCG sc animals along PC1, with activation of CD8+ T cells and their recruitment into the lung parenchyma as well as the upregulation of IFN-inducible markers by myeloid cells as the driving factors (**Fig5B and Supplementary Table 1**). In contrast, BCG iv animals clustered together irrespective of SARS-CoV-2 challenge. PC1 and PC2 separated the BCG iv group from PBS iv and BCG sc animals, with PC2 representing variables uniquely driven by iv BCG inoculation (**Fig5A**). The latter variables included increased numbers of CD4+ T cells and cDC1, reduced numbers of NK cells and the induction of iNOS in myeloid cells (**Supplementary Table 1 and Fig5B**). Hierarchal clustering of cytokine, chemokine and flow cytometry data highlighted the unique signatures present in animals that were iv BCG inoculated then challenged with SARS-CoV-2, as well as the minimal changes observed between WT and transgenic animals within this treatment group (**Fig5B**).

**Fig 5:**
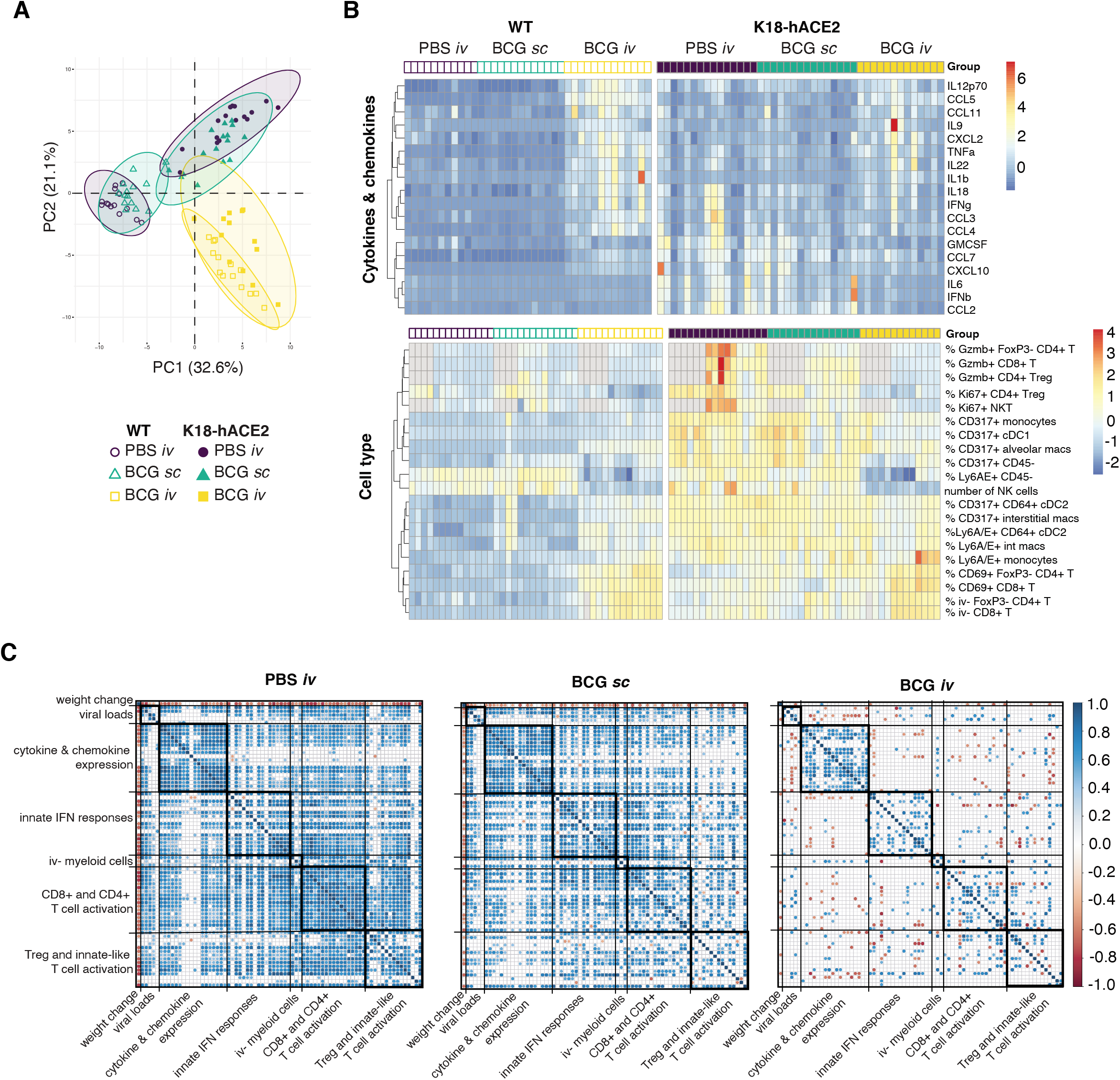
Prior *iv* BCG inoculation suppresses inflammatory responses following SARS-CoV-2 infection: lack of correlation with viral load. **(A-C)** K18-hACE2 mice or non-transgenic littermate controls (WT) were inoculated with 10^6^ CFU BCG Pasteur by subcutaneous (*sc*) or intravenous (*iv*) routes. Control animals received the same volume of PBS *iv*. At 42 days post BCG administration, mice were infected with 10^3^ TCID_50_ SARS-CoV-2 (WA1/2020) by intranasal instillation. Tissues were harvested for viral titers, flow cytometry analysis and cytokine measurements 5 days after viral challenge. **(A**) Principal component analysis (PCA) incorporating weight change, viral copies in nasal turbinates, lungs and brain, cytokine levels in lung homogenate and cell type frequencies and numbers in lung single-cell suspensions. The full data set is shown in Supplementary Table 1. (**B)** Hierarchical clustering of the top 10 cytokines and cell types that contribute to variation along the first and second components of the PCA is represented as a heatmap. Experimental groups are indicated above the heatmap. Gray boxes denote data not collected. (**C)** Spearman’s correlation analyses of all parameters utilized in the PCA were performed for the PBS *iv*, BCG *sc* and BCG *iv* groups separately. Positive and negative correlations are shown in blue and red as represented in the key and only statistically significant correlations (adjusted p-value <0.05) are indicated.

We next performed Spearman’s correlation analyses to assess the relationship between all parameters measured in our study. To do so, a pairwise comparison for each of the variables (e.g. viral titer, cytokine level, cell number) from the individual virus-challenged WT and K18-hACE2 mice was performed independently for the PBS iv, BCG sc and BCG iv groups. Any significant correlations observed would therefore be a readout of a SARS-CoV-2-induced response. Strong positive correlations between variables were observed in animals that received PBS iv. The notable exception was weight change which, as expected, was negatively associated with many of the variables measured. Viral loads positively correlated with induction of pro-inflammatory cytokines and chemokines, activation and IFN priming of myeloid cells, as well as cytotoxic activity by lymphocytes and their recruitment into the lung parenchyma (**Fig5C**). Cytokine, chemokine and cellular response modules that are co-expressed after SARS-CoV-2 challenge were also identified from this analysis (**Fig5C and Supplementary Table 2**). For example, markers of CD4+ and CD8+ T cell activation, such as PD1, CD69, CD44 and Gzmb, displayed strong positive correlations. Similar modules and correlations were observed in animals that received BCG sc, although these were less obvious than seen with PBS iv controls, particularly for cellular responses. The majority of these significant correlations were strikingly absent from the cohort of animals that received BCG iv prior to challenge, and importantly the associations with viral loads.

Collectively our findings suggest that while iv BCG does reduce viral burden, this alone may not be sufficient to explain the pronounced inhibition of the SARS-CoV-2 inflammatory response and protection from lethal challenge observed in these animals. Instead, we propose that a major effect of prior iv BCG exposure is to limit the pathological effects of the innate response to the virus and speculate that this outcome stems from the local effects of BCG-induced IFNγ on the pulmonary epithelioid and myeloid compartments. While iv administration of BCG is currently not a clinically acceptable practice, the experimental proof-of-concept that prior iv BCG can trigger potent protection against lethal SARS-CoV-2 challenge may be of value in the design of other strategies for COVID-19 prophylaxis that target the innate response to the virus.

## Methods

### Mice

Female, 7–9-week-old, B6.Cg-Tg(K18-ACE2)2Prlmn/J hemizygous and non-carrier control animals (JAX34860) were purchased from Jackson Laboratories (Bar Harbor, ME). Female 7-9-week-old B6 mice were acquired from the NIAID Contract Facility at Taconic Farms. Mice were housed under specific pathogen–free conditions with *ad libitum* access to food and water and were randomly assigned to experimental groups. All animal studies were performed in accordance with institutional guidelines and were conducted in Assessment and Accreditation of Laboratory Animal Care–accredited Biosafety Level 2 and 3 facilities at the NIAID/National Institutes of Health (NIH) using a protocol (LPD-99E) approved by the NIAID Animal Care and Use Committee.

### Virology

The SARS-CoV-2 WA1/2020 strain (Pango lineage A), originally isolated at the Center for Disease Control and Prevention (Atlanta, GA) and representative of SARS-CoV-2 viruses circulating early during the pandemic, was obtained from BEI Resources and propagated in tissue culture in Vero CCL81 cells (ATCC). The SARS-CoV-2 USA/CA_CDC_5574/2020 isolate, an alpha variant of concern (Pango lineage B.1.1.7), was obtained from BEI resources and propagated in Vero cells overexpressing TMPRSS2, kindly provided by Dr Jonathan Yewdell (NIAID). Vero cells were maintained in DMEM medium supplemented with glutamax and 10% FBS. Vero-TMPRSS2 cells were maintained in were maintained in DMEM medium supplemented with glutamax, 10% FBS and 250μg/ml Hygromycin B gold (InvivoGen). Virus stock production was performed under BSL-3 conditions by the NIAID SARS-CoV-2 Virology Core using DMEM medium supplemented with glutamax and 2% FBS. At 48h post inoculation, culture supernatant and cells were collected, clarified by centrifugation for 10 min at 4°C. Supernatant was collected, aliquoted and frozen at −80°C. Virus titers were determined by TCID_50_ assay in Vero E6 cells (ATCC CRL-1586) using the Reed and Muench calculation method. Full genome sequencing was performed at the NIAID Genomic Core (Hamilton, MT). The WA isolate stock used in this study was 6 passages from isolation and contained 4 single nucleotide polymorphisms compared to the reference sequence (MN985325.1): C23525T, C26261T, C26542T and T28853A.

### BCG

BCG Pasteur was propagated in 7H9 broth supplemented with OADC until mid-log phase. Bacteria were harvested, washed thrice and frozen down in aliquots until use. Colony forming units were enumerated by culturing on 7H11 agar for 3 weeks at 37°C.

### Vaccinations and infections

BCG vaccinations were performed by administering 10^6^ CFU in PBS containing 0.05% Tween-80 by subcutaneous or intravenous injection. Control animals received the same volume of PBS with 0.05% Tween80 by intravenous injection.

SARS-CoV-2 infections were performed under BSL3 containment. Animals were anesthetized by isoflurane inhalation and 10^3^ TCID50 was administered by intranasal instillation. Following infection, mice were monitored daily for weight change and clinical signs of disease by a blinded observer who assigned each animal a disease score based on the following criteria: 0) no observable signs of disease; 1) hunched posture, ruffled fur and/or pale mucous membranes; 2) hunched posture and ruffled fur with lethargy but responsive to stimulation, rapid/shallow breathing, dehydration; 3) moribund.

### Determination of viral copies and ACE2 expression by quantitative PCR

Lung, brain and nasal turbinates were homogenized in Trizol and RNA was extracted using the Direct-zol RNA Miniprep kit following the manufacturer’s instructions. E gene gRNA was detected using the QuantiNova Probe RT-PCR Kit and protocol and primers (forward primer: 5’-ACAGGTACGTTAATAGTTAATAGCGT-3’, reverse primer:5’-ATATTGCAGCAGTACGCACACA-3’) and probe (5’-FAM-ACACTAGCCATCCTTACTGCGCTTCG-3IABkFQ-3’) as previously described (Corman et al., 2020). The standard curve for each PCR run was generated using the inactivated SARS-CoV2 RNA obtained from BEI (NR-52347) to calculate the viral copy number in the samples. Human ACE2 transgene expression in the lung was quantified using the TaqMan probe (Assay ID: Hs01085333_m1 FAM) and Fast Advanced Master Mix following manufacturer’s instructions. The expression of mouse GAPDH gene (Assay ID: Mm99999915_g1 VIC) was used as internal control and the relative expression of hACE2 was calculated using the 2^−ΔΔCT^ method. Identical lung and brain portions were utilized for all experiments to generate comparable results.

### Determination of viral titers by TCID50 assay

Viral titers from lung and brain homogenate were determined by plating in triplicate on Vero E6 cells using 10-fold serial dilutions. Plates were stained with crystal violet after 96 hours to assess cytopathic effect (CPE). Viral titers were determined using the Reed-Muench method.

### Preparation of single cell suspensions from lungs

Lung lobes were diced into small pieces and incubated in RPMI containing 0.33mg/mL Liberase TL and 0.1mg/mL DNase I (both from Sigma Aldrich) at 37°C for 45 minutes under agitation (200rpm). Enzymatic activity was stopped by adding FCS. Digested lung was filtered through a 70μm cell strainer and washed with RPMI. Red blood cells were lysed with the addition of ammonium-chloride-potassium buffer (Gibco) for 5 minutes at room temperature. Cells were then washed with RPMI supplemented with 10% FCS. Live cell numbers were enumerated using AOPI staining on a Cellometer Auto 2000 Cell Counter (Nexcelom).

For assessment of cytokine production, single cell suspensions were incubated in FCS-supplemented RPMI in the presence of 1X Protein Transport Inhibitor Cocktail (eBioscience) for 5 hours.

### Flow cytometry

To label cells within the pulmonary vasculature for flow cytometric analysis, 2μg anti-CD45 (30-F11; Invitrogen) was administered by intravenous injection 3 minutes prior to euthanasia.

Single-cell suspensions prepared from lungs were washed twice with PBS prior to incubating with Zombie UV™ Fixable Viability Dye and TruStain FcX™ (clone 93; both from BioLegend) for 15 minutes at room temperature. Cocktails of fluorescently-conjugated antibodies diluted in PBS containing 2mM EDTA, 0.01% sodium azide, 2% FCS and 10% Brilliant Stain Buffer (BD) were then added directly to cells and incubated for a further 20 minutes at room temperature. Anti-CD4 (GK1.5), anti-CD11b (M1/70) and anti-CD26 (H194-112) were from BD OptiBuild. Anti-CD19 (GL3), anti-CD45 (30-F11), anti-Siglec F (E50-2440), anti-TCR-beta chain (H57-597) and anti-TCR-gamma-delta (GL3) were from BD Horizon. Anti-CD8-alpha (53-6.7) was from Invitrogen. Anti-CD11c (N418), anti-CD19 (6D5), anti-CD44 (IM7), anti-CD64 (X54-5/7.1), anti-CD69 (H1.2F3), anti-CD86 (GL-1), anti-CD88 (20/70), anti-CD279 (PD1, clone 29F. 1A12), anti-CD317 (BST2, clone 927), anti-IA/IE (MHCII, clone M5/114), anti-NK1.1 (PK136), anti-Ly6A/E (Sca-1, clone D7), anti-Ly6C (HK1.4), anti-Ly6G (1A8), anti-TCR-beta chain (H57-597), anti-TCR-gamma-delta (GL3) and anti-XCR1 (ZET) were from BioLegend. SARS-CoV-2 S(539-546) and N(119-227) tetramers were from the NIH Tetramer Core.

Cells were incubated in eBioscience™ Transcription Factor Fixation and Permeabilization solution (Invitrogen) for 2-18 hours at 4°C and stained with cocktails of fluorescently-labeled antibodies against intracellular antigens diluted in Permeabilization Buffer (Invitrogen) for 20 minutes at room temperature followed by 20 minutes at 4°C. Anti-Granzyme B (GB11) and anti-Ki67 (B56) were from BD Pharmingen. Anti-FoxP3 (FJK-16s), anti-NOS2 (CXNFT) and anti-Tbet (4B10) were from Invitrogen. Anti-IL-6 (MP5-20F3) and anti-TNF-alpha (MP6-XT22) were from BioLegend.

Compensation was set in each experiment using UltraComp eBeads™ (Invitrogen) and dead cells and doublets were excluded from analysis. All samples were collected on a FACSymphony A5 SORP™ flow cytometer (BD) and analyzed using FlowJo software (version 10, BD). A minimum of 1000 events were analyzed per gated population.

### Multiplex Cytokine Array

Cytokines were assessed in lung homogenate using a ProcartaPlex Luminex kit (ThermoFisher) according to the manufacturers’ instructions and measured using a MagPix Instrument (R&D Systems). Total protein was measured by Pierce™ Protein Assay (ThermoFisher). Cytokine levels were standardized to total protein content.

### Histology

Tissues were fixed in 10% neutral buffered formalin for 48-72 hours and transferred into 30% sucrose, and embedded in paraffin. Embedded tissues were sectioned at 5 um and dried overnight at 42°C prior to staining. Specific anti-CoV immunoreactivity was detected using a SARS-CoV-2 nucleoprotein antibody (Genscript) at a 1:1000 dilution. The secondary antibody was the Vector Laboratories ImPress VR antirabbit IgG polymer (cat# MP-6401). The tissues were then processed for immunohistochemistry using the Discovery Ultra automated stainer (Ventana Medical Systems) with a ChromoMap DAB kit (Roche Tissue Diagnostics cat#760–159). All tissue slides were evaluated by a board-certified veterinary pathologist.

### Statistical analyses

P-values were determined by Student’s unpaired *t*-test or Mann-Whitney test when comparing two groups, or by One-Way ANOVA with Tukey’s post-test or Kruskal-Wallis test with Dunn’s post-test when comparing three or more groups using GraphPad Prism software (v9). P-values below 0.05 were considered statistically significant.

Weight change, viral copies, multiplex cytokine and flow cytometry measurements generated from WT and K18-hACE2 mice for the three experimental groups (PBS *iv*, BCG *sc* and BCG *iv*) from three independent experiments were compiled and principal component analysis (PCA) was performed and visualized in R version 4.1.0 with packages *FactoMineR* and *factoextra*. The top 10 variables contributing to the first two principal components were identified for the cytokine and flow cytometry data and these measurements were normalized, clustered and visualized as a heatmap using the R package *pheatmap*. Pairwise Spearman’s correlation values were calculated for the same read-outs used for PCA for each of the three experimental groups. Correlations that were found to be significant following Benjamini-Hochberg correction for multiple testing were visualized using the R packages *RcmdrMisc and corrplot*.

### Figure visualization

Figures were generated in Adobe Illustrator and R incorporating images from Biorender.com.

## Author contributions

Conceptualization: KLH, SN, AS. Methodology: KLH, SN, CSC, SJR, PJB. Investigation: KLH, SN, CSC, DO, SDO, PJB, EC. Resources: NLG, BAPL, RJ, SMB. Data curation and analysis: KLH, SN, CSC. Writing – original draft: KLH, AS. Writing – review and editing: All Authors. Visualization: KLH, SN, CSC. Supervision: FR, KDM-B, SMB, AS. Funding acquisition: AS.

## Acknowledgements

We are grateful to Drs. Dragana Jankovic, Daniel Barber, Eduardo Amaral, Christine Nelson, Patricia Earl, Jeffrey Americo and Heather Hickman for critical discussion. We also thank Virgilio Bundoc and Nickiana Lora for technical assistance; Dr. Craig Martens and the RML Genomics Unit for viral sequencing; the NIAID Research Technologies Branch for assistance with flow cytometry; the NIAID animal care staff; and the NIH Tetramer Core. KLH was partially supported by a Malaghan Institute Postdoctoral Fellowship. This research was funded by the Intramural Program of NIAID, NIH.

