## Supplemental Figures for "Intravenous administration of BCG protects mice against lethal SARS-CoV-2 challenge"

Hilligan, Namasivayam *et al.*

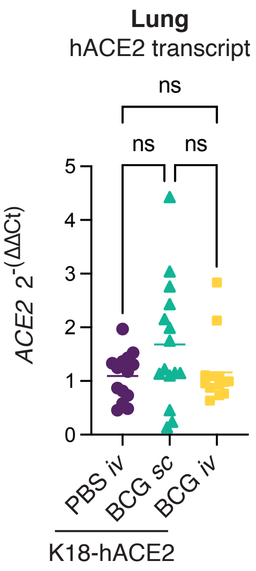

Lung
hACE2 transcript
ns
ns
ns
ACE2 2^-(ΔΔCt)
0
1
2
3
4
5
PBS iv
BCG sc
BCG iv
K18-hACE2

**Supplementary Fig 1: Prior *iv* BCG inoculation does not impact expression of hACE2.**

K18-hACE2 mice were inoculated with $10^6$ CFU BCG Pasteur by subcutaneous (*sc*) or intravenous (*iv*) injection. Control animals received the same volume of PBS *iv*. At 42 days post BCG administration, mice were infected with $10^3$ $TCID_{50}$ SARS-CoV-2 (WA1/2020) by intranasal instillation. Lungs were collected 5 days after viral challenge and assessed for expression of human ACE2 by RT-qPCR. Statistical significance was assessed by One-Way ANOVA with Tukey post-test. ns $p>0.05$. Data are pooled from 3 independent experiments each with 4-5 mice per group.

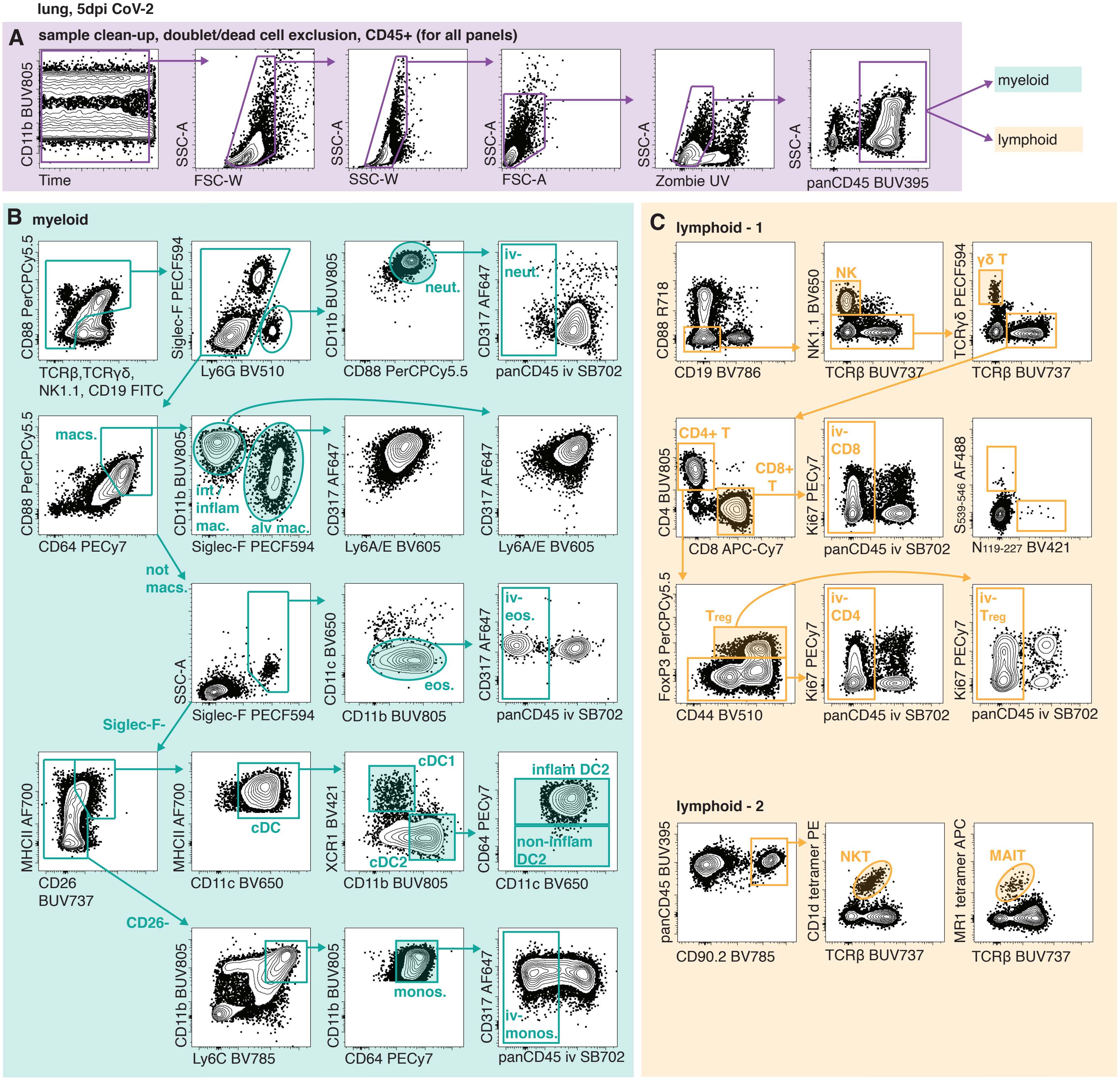
lung, 5dpi CoV-2
A
sample clean-up, doublet/dead cell exclusion, CD45+ (for all panels)
CD11b BUV805
Time
SSC-A
FSC-W
SSC-A
SSC-W
SSC-A
FSC-A
SSC-A
Zombie UV
SSC-A
panCD45 BUV395
myeloid
lymphoid
B
myeloid
CD88 PerCPCy5.5
TCRβ,TCRγδ, NK1.1, CD19 FITC
Siglec-F PECF594
Ly6G BV510
CD11b BUV805
CD88 PerCPCy5.5
neut.
CD317 AF647
panCD45 iv SB702
iv-neut.
CD88 PerCPCy5.5
CD64 PECy7
macs.
CD11b BUV805
Siglec-F PECF594
int / inflam mac.
alv mac.
CD317 AF647
Ly6A/E BV605
CD317 AF647
Ly6A/E BV605
not macs.
SSC-A
Siglec-F PECF594
CD11c BV650
CD11b BUV805
eos.
CD317 AF647
panCD45 iv SB702
iv-eos.
Siglec-F-
MHCII AF700
CD26 BUV737
MHCII AF700
CD11c BV650
cDC
XCR1 BV421
CD11b BUV805
cDC1
cDC2
CD64 PECy7
CD11c BV650
inflam DC2
non-inflam DC2
CD26-
CD11b BUV805
Ly6C BV785
CD11b BUV805
CD64 PECy7
monos.
CD317 AF647
panCD45 iv SB702
iv-monos.
C
lymphoid - 1
CD88 R718
CD19 BV786
NK1.1 BV650
TCRβ BUV737
NK
TCRγδ PECF594
TCRβ BUV737
γδ T
CD4 BUV805
CD8 APC-Cy7
CD4+ T
CD8+ T
Ki67 PECy7
panCD45 iv SB702
iv-CD8
S539-546 AF488
N119-227 BV421
FoxP3 PerCPCy5.5
CD44 BV510
Treg
Ki67 PECy7
panCD45 iv SB702
iv-CD4
Ki67 PECy7
panCD45 iv SB702
iv-Treg
lymphoid - 2
panCD45 BUV395
CD90.2 BV785
CD1d tetramer PE
TCRβ BUV737
NKT
MR1 tetramer APC
TCRβ BUV737
MAIT

**Supplementary Fig 2: Gating strategies for identifying cell types by flow cytometry.**

**(A-C)** Single cell suspensions were prepared from the lungs of animals intravenously injected with a fluorescently labeled panCD45 antibody 3 minutes prior to euthanasia to enable identification of cells located within the pulmonary vasculature. Cell suspensions were stained with cocktails of fluorescent antibodies and analyzed by flow cytometry. **(A)** Gating strategy to identify CD45+, live, singlets. **(B)** Gating strategy to identify myeloid subsets by flow cytometry. **(C)** Gating strategies to identify lymphoid subsets by flow cytometry.

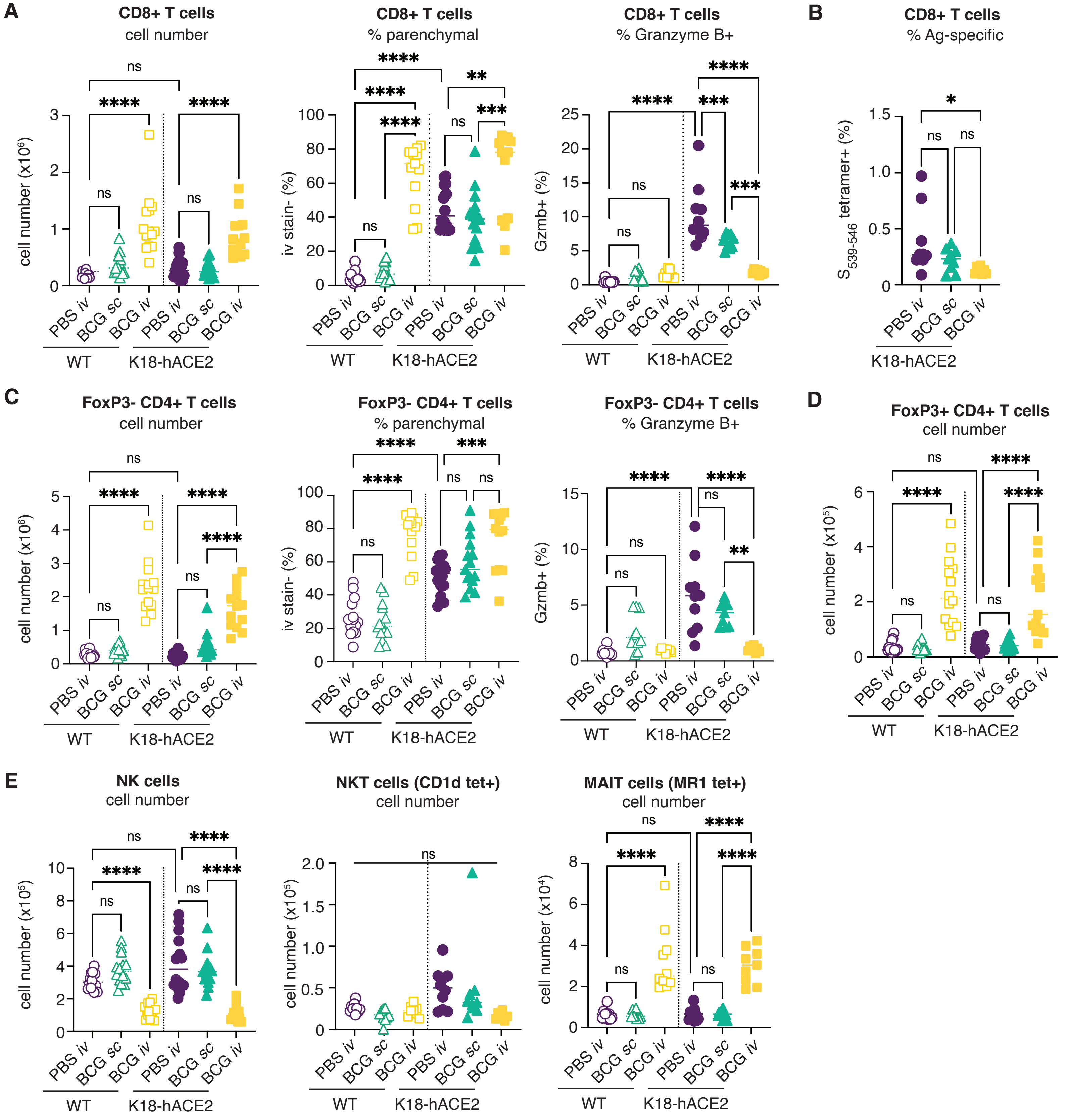
A
CD8+ T cells
cell number
cell number (x10^6)
CD8+ T cells
% parenchymal
iv stain- (%)
CD8+ T cells
% Granzyme B+
Gzmb+ (%)
B
CD8+ T cells
% Ag-specific
S539-546 tetramer+ (%)
C
FoxP3- CD4+ T cells
cell number
cell number (x10^6)
FoxP3- CD4+ T cells
% parenchymal
iv stain- (%)
FoxP3- CD4+ T cells
% Granzyme B+
Gzmb+ (%)
D
FoxP3+ CD4+ T cells
cell number
cell number (x10^5)
E
NK cells
cell number
cell number (x10^5)
NKT cells (CD1d tet+)
cell number
cell number (x10^5)
MAIT cells (MR1 tet+)
cell number
cell number (x10^4)
PBS iv
BCG sc
BCG iv
WT
K18-hACE2
ns
*
**
***
****

**Supplementary Fig 3: Prior *iv* BCG administration limits bystander cytotoxic responses in SARS-CoV-2 challenged mice.**

**(A-E)** K18-hACE2 mice or non-transgenic littermate controls (WT) were inoculated with $10^6$ CFU BCG Pasteur by subcutaneous (*sc*) or intravenous (*iv*) routes. Control animals received the same volume of PBS iv. At 42 days post BCG administration, mice were infected with $10^3$ $TCID_{50}$ SARS-CoV-2 (WA1/2020) by intranasal instillation. Lungs were collected 5 days after viral challenge and assessed by flow cytometry. Gating strategy is shown in Supplementary Fig 2. **(A)** Number of CD8+ T cells, frequency of CD8+ T cells negative for iv stain and frequency of CD8+ T cells expressing Granzyme B. **(B)** Frequency of CD8+ T cells positive for the S539-546 tetramer. **(C)** Number of FoxP3- CD4+ T cells, frequency of FoxP3- CD4+ T cells negative for iv stain and frequency of FoxP3- CD4+ T cells expressing Granzyme B. **(D)** Number of FoxP3+ CD4+ T cells. **(E)** Number of NK, NKT and MAIT cells. Statistical significance was assessed by One-Way ANOVA with Tukey post-test. ns $p>0.05$; *$p<0.05$; **$p<0.01$; ***$p<0.001$, ****$p<0.0001$. Data are pooled from 3 independent experiments each with 4-5 mice per group.
